## Supplementary figures and images for "African eggplant-associated virus: characterization of a novel tobamovirus identified from *Solanum macrocarpon* and assessment of its potential impact on tomato and pepper crops"

### S1 Figure

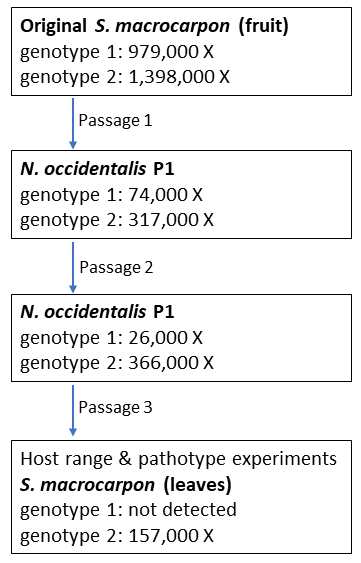

### S2 Figure

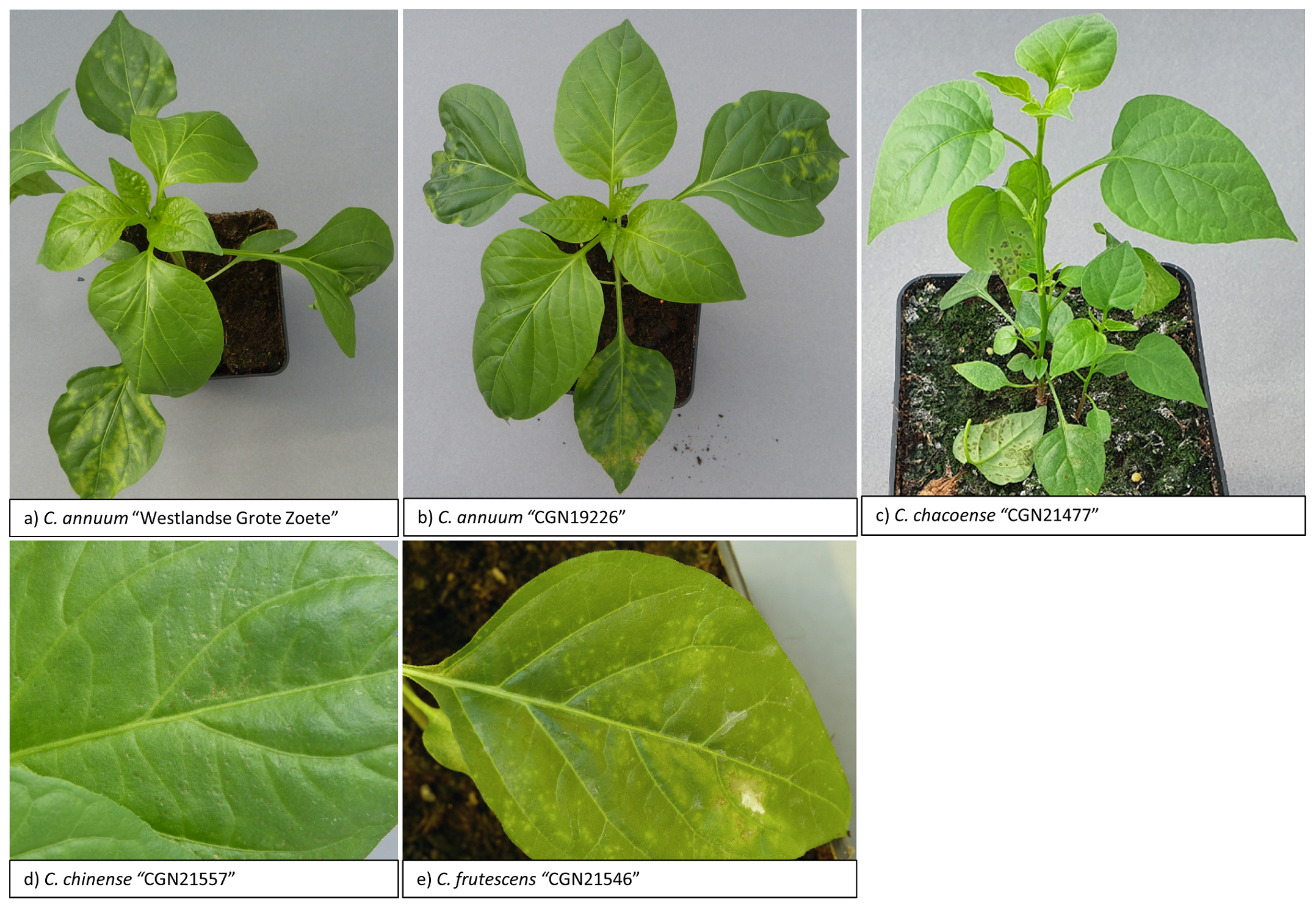
